# Mechanism of glucocorticoid receptor activation regulated expression of thrombospondin-1

**DOI:** 10.1101/2023.04.13.536820

**Authors:** Yan-Yan Wang, Yi-Yi Song, Wen-Yi Jiang, Hao-Tian Zhang, Jing-Wei Chen, Koji Murao, Wan-Ping Sun, Guo-Xing Zhang

## Abstract

**Objective:** Thrombospondin-1 (TSP-1) plays an important role in platelet activation and aggregation and aggravates thrombosis. Chronic stress can cause a variety of diseases, including coagulation disorders, increased thrombosis, atherosclerosis, and a series of cardiovascular and cerebrovascular diseases. However, it is still unknown how chronic stress regulates the expression of TSP-1 after glucocorticoid receptor activation.

**Approach and Results:** rats chronic unpredictable mild stress model was applied and the changes of TSP-1 and microRNAs in plasma were examined. Effects of glucocorticoid receptor activation on human umbilical vein endothelial cells and platelets were observed. Glucocorticoid receptor (GR) activation upregulated the expression of TSP-1 and downregulated the expression of microRNA-1-3p accompanied with increase of phosphorylation of p38 mitogen-activated protein kinase (MAPK) and argonaute-2 (AGO-2). Blockade of p38 MAPK phosphorylation resulted in decrease of phosphorylation level of AGO-2, increase of microRNA-1-3p expression, and decrease of TSP-1 expression. Transfection of AGO-2 Y393F point mutant plasmid, increased microRNA-1-3p expression and decreased TSP-1 expression, transfection of microRNA-1-3p mimic also decreased TSP-1 expression, while transfection of microRNA-1-3p inhibitor increased TSP-1 expression. Finally, GR activation led to an increase in the phosphorylation level of p38 MAPK in platelets and an increase in the level of TSP-1 in the supernatant.

**Conclusions:** our study demonstrates that GR activation in HUVEC stimulates the phosphorylation of p38 MAPK, which in turn promotes the phosphorylation of AGO-2 and inhibits the maturation of microRNA-1-3p, leading to elevated expression of TSP-1, GR activation in platelets leads to the release of TSP-1.

Graphical Abstract
HSS: Hydrocortisone sodium succinate

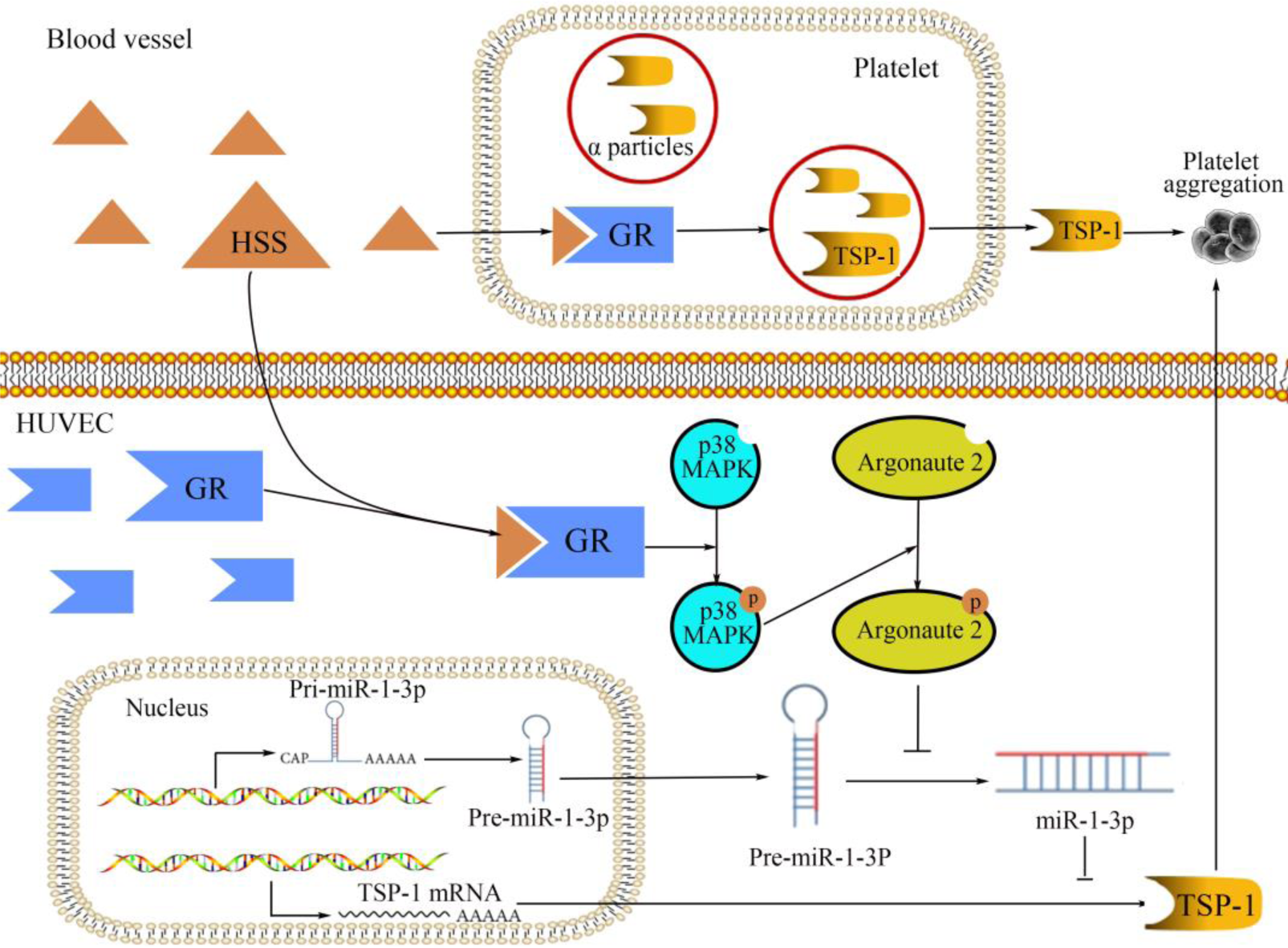

## Highlights

This study revealed the mechanism by which GR activation regulates TSP-1 MicroRNA-1-3p targets the regulation of TSP-1 expression.

Glucocorticoid receptor activation promotes the phosphorylation of p38 MAPK and then promotes the phosphorylation of AGO-2, regulates the expression of microRNA-1-3p.

GR activation in platelets promotes the release of TSP-1.

## 1. Introduction

Stress is a non-specific and complex response of a human body submitted to a stressor, which response to an adaptive function. Stressors included physiological and psychological stressors(1). Psychological stress refers to the process of changes in physiological and psychological functions that are produced autonomously by the organism in response to the stimulus of a stressor. Physiological stress refers to the process of functional changes in the endocrine system and metabolic system after an organism is stimulated. Stress can be divided into acute or chronic stress. Acute stress usually refers to the short-term adaptation of the stress response system to stressors in an emergency, such as an increase in blood glucose, blood pressure, and heart rate(2). At the same time, the body temporarily produces catecholamines and corticosteroids that improve responsiveness and mobility. Therefore, acute stress is often conducive to the body’s adaptation to adverse environments. Chronic stress is considered to be the result of cumulative, repetitive, or prolonged exposure to stressors, including occupational stress, social isolation, and unusual adversity(3). Chronic stress can suppress adaptive immunity and thus predispose the body to disease. It has been shown that chronic stress can lead to a variety of diseases, such as insomnia, and gastrointestinal disorders^(4, 5)^, mental illness, cancer(6), and a range of cardiovascular diseases^(7, 8)^. Chronic stress can modulate vascular endothelial cell function, increase thrombosis, dys-coagulation, increased blood clots, and atherosclerosis, all of which may trigger arrhythmias and myocardial infarction((9–12)). Our previous study also confirmed that stress can exacerbate deep venous thrombosis (DVT)(13), but the underlying mechanisms need to be further investigated.

Thrombospondin-1 (TSP-1), a multifunctional glycoprotein composed of homotrimers, was first found in platelets(14), and it is expressed in a variety of cellular tissues including endothelial cells, macrophages, monocytes, and smooth muscle cells(15). It mediates a variety of biological processes by interacting with a variety of receptors, some cytokines, and other matrix proteins^(16, 17)^. TSP-1 can interact with CD36 to inhibit angiogenesis(18), it can inhibit the proliferation of human microvascular endothelial cells by interacting with very low density lipoprotein receptor (VLDLR)(19), and it can interact with apolipoprotein E receptor 2 (APOER2) to promote neuronal migration(20). TSP-1 can affect cell adhesion, migration, and regulate angiogenesis(18). Recent studies have shown that TSP-1 can bind to its receptor CD47 and antagonize the NO/cGMP pathway, thereby inducing platelet aggregation^(21, 22)^, it regulate thrombus adhesion by protecting vWF from ADAMTS cleavage(23), induce platelet activation, and promote hemostasis^(24, 25)^. So, does TSP-1 plays an important role in the exacerbation of thrombosis caused by chronic stress?

MicroRNA is a class of highly conserved endogenous non-coding RNA of 21-25 nucleotides in length(26). It usually plays a role in post-transcriptional regulation by binding to the 3’untranslated region (3’UTR) of the target gene and destabilizing or translationally silencing the mRNA to regulate the target gene(27). It has been shown that stress affects microRNA expression(28), and chronic stress leads to significant downregulation of miR-211-5p in the rat hippocampus(29). Microarray analysis of myocardial tissues from rats subjected to chronic stress revealed changes in 55 microRNAs, 30 of which were down-regulated and 25 were up-regulated(30). In addition, the expression of several hemostatic factors is regulated by microRNAs^(31, 32)^, including key procoagulants and inhibitors of the coagulation process((33–35)). Chronic stress can affect microRNA maturation through p38 MAPK-mediated phosphorylation of argonaute-2 (AGO-2)((36–38)).

AGO-2 is one of four members of the argonaute protein family in mammals^(39, 40)^, AGO-2 plays an important role in RNA-induced silencing complex (RISC), in which microRNA are loaded on AGO-2 and complementary to the mRNA of the target gene to inhibit the translation of the target gene^(41, 42)^. AGO-2 is a key regulator of microRNA processing and it has endonuclease activity(43), which can bind to Dicer(44) and mediate microRNA maturation through endonuclease activity^(45, 46)^, previous studies have shown that the increased phosphorylation of AGO-2 Y393 blocks the binding of AGO-2 to Dicer and inhibits microRNA maturation(47). In addition, AGO-2 can maintain the stability of microRNA and protect microRNA from degradation((48–50)).

Chronic stress releases cortisol (humans) and corticosterone (rodents) leading to glucocorticoid receptor activation, in present study, our results showed that glucocorticoid receptor activation in vascular endothelial cells stimulated the phosphorylation of p38 MAPK, which in turn promoted the phosphorylation of AGO-2 and inhibited the maturation of microRNA-1-3p, leading to the increased expression of TSP-1. Glucocorticoid receptor activation in platelets leads to the release of TSP-1 and increases the phosphorylation of p38 MAPK.

## 2. Materials and methods

### 2.1 Cell Culture and Reagents

Human umbilical vein endothelial cells (HUVECs) (gifts from Hematology Institute of Soochow University). The cells were cultured in Dulbecco’s modified Eagle’s medium (DMEM; Gibco) supplemented with 10% fetal bovine serum (FBS; Gibco), 100 μg/mL streptomycin, and 100 units/mL penicillin, 0.01 mg/mL heparin sodium (Wanbang Pharmaceutical) at 37 °C in a humidified atmosphere of 5% CO_2_. Hydrocortisone sodium succinate was obtained from Tianjin Biochemical Pharmaceutical Co. RU486, corticosterone was purchased from Sigma. BIRB796 was purchased from Selleckchem.

### 2.2 Animals and groups

Male Sprague-Dawley rats (ten-week-old) used in this study were obtained from Shanghai Laboratory Animal Center. All rats were raised in standard housing conditions (22±1℃ with light/dark cycle for 12 hours) without restriction of food and water. All rats were divided into two groups: control (Con), chronic unpredictable mild stress (CUMS). All the protocols were conformed to the European Convention for the Protection of Vertebrate Animals used for Experimental and other Scientific Purposes. All experiments were approved by the Committee on Animal Resources of Soochow University.

### 2.3 Chronic unpredictable mild stress (CUMS) procedure

The CUMS procedure was performed as described(51) with a slight modification. The animals in CUMS groups were exposed to unpredicted chronic mild stressors for 4 weeks. Rats in CUMS group were subjected to different stressors: fast for 24 hours, water deprivation for 24 hours, restraint for 8 hours, Light/dark alternately for 3 hours (alternate every 30min), cold swimming for 5 min (at 4 °C), level shaking for 30 min, cage tilting 45°and wet bedding for 24 hours, inversion of the light/dark cycle for 24 hours, rats were isolated in a single cage for 3 hours, all rats were housed in a crowd cage for 3 hours, tail nip for 1 min (1 cm from the end of the tail). Two or three stressors were randomly selected each day, and the same stressors were not applied consecutively for two days, so animals could not predict the occurrence of stimulation. The control group was undisturbed except for necessary procedures such as routine cage cleaning.

### 2.4 Measurement of plasma PAF, TSP-1 and VWF levels

After the rats were anesthetized with sodium pentobarbital (45 mg/kg i.p.), blood was collected from the abdominal aorta. Plasma concentrations of platelet activated factor (PAF), TSP-1 and von Willebrand factor (VWF) were determined using commercially available enzyme-linked immunosorbent assays (ELISA) (ELISA kit for Platelet Activating Factor (PAF), USCN Wuhan, China), (rat thrombospondin (THBS1) ELISA kit, USCN Wuhan, China), (rat vWF ELISA kit, Xiamen Lunchangshuo Biotechnology Co., Ltd., Xiamen, China).

### 2.5 Platelet extraction and culture

Blood was obtained from the abdominal aorta after the rats were anesthetized with sodium pentobarbital, put it into a centrifuge tube containing 7.6% sodium citrate, afterward, platelet-rich plasma (PRP) was obtained after centrifugation at 1000 rpm/min for 10 min. Transfer the supernatant to another centrifuge tube, centrifuge at 3000rpm for 10min, discard the supernatant, add platelet buffer (137 mM NaCl, 2.7 mM KCl, 2 mM MgSO4, 10 mM HEPES, 5 mM glucose, pH 7.4) without calcium to wash twice, centrifuge at 3000rpm for 10min, discard the supernatant, and the precipitate is platelets. The platelets were cultured in Dulbecco’s modified Eagle’s medium (DMEM; Gibco) supplemented with 100 μg/mL streptomycin, and 100 units/mL penicillin, at 37 °C in a humidified atmosphere of 5% CO2.

### 2.6 Immunofluorescence

Cells were first washed with phosphate-buffered saline (PBS) and fixed with 4% PFA for 20 min. Add immunostaining permeabilizer Tritenx-100 (P0096, Beyotime, Shanghai, China) to permeabilize, and block with immunostaining blocking solution (P0102, Beyotime, Shanghai, China) for 1 hours. The slides were then incubated with primary antibodies overnight (4°C) and, subsequently, with a secondary antibody for 2 hours (at room temperature). The primary antibody used in this study was anti-TSP-1 (1:100, 37879, Cell Signaling Technology) and the secondary antibody was Alexa Fluor 594-conjugated Affinipure goat anti-rabbit igG(H+L). The nucleus was counterstained with DAPI for 5 minutes, Confocal images were taken using a confocal laser-scanning microscope (Olympus FV1200, Japan).

### 2.7 RNA Preparations and Quantitative Real-Time PCR

RNA was extracted and collected by using TRIzol solvent. Then reverse transcription reaction was conducted by means of TaKaRa reverse transcription reagents (TaKaRa, Dalian, China). The quantitative real-time polymerase chain reaction was performed using an SYBR Premix. GAPDH was used as a reference gene for the normalization of the relative qRT-PCR data. Data were calculated by the ΔΔCt method. The sequences of the primers were as follows: GAPDH-F, 5’-GAC GCT GGT GCT GGT ATT GCT-3’, GAPDH-R, 5’-CTA CTC CTT GGA GGC CAT GTG T-3’. TSP-1-F, 5’-TTT GAC ATC TTT GAA CTC ACC G-3’, TSP-1-R, 5’-AGA AGG AGG AAA CCC TTT TCT G-3’.

Plasma miRNA was extracted using Serum/Plasma miRNA Extraction Kit (HaiGene, Harbin, China), and reverse transcription was performed using TaqMan miRNA cDNA Synthesis Kit (Serum/Plasma) (HaiGene, Harbin, China), miRNA in cells was extracted using miRNA Extraction Kit (Tissue & Cell) (HaiGene, Harbin, China), reverse transcription using HG TaqMan miRNA cDNA Synthesis Kit (HaiGene, Harbin, China), Quantitative real-time polymerase chain reaction used HG TaqMan miRNA qPCR Kit (HaiGene, Harbin, China), miR-16 was used as an internal reference for miRNAs in plasma, and U6 was used as an internal reference for miRNAs in cells. Data were calculated by the ΔΔCt method.

### 2.8 Western blot

Cell lysates were prepared in RIPA buffer (Beyotime), and cell lysates containing fresh protease and phosphatase inhibitors (roche) and PMSF (Solarbio) (1 mM) were added to cells and placed on ice for 30 min, and were quantified using a BCA Protein Assay Kit (Beyotime). Equal amounts of protein were loaded and electrophoresed in SDS-PAGE and transferred to PVDF membranes to be probed by primary antibodies (TSP-1, 1:1000, 37879, CST; phospho-p38, 1:1000, 4511, CST; p38 MAPK, 1:1000, 8690, CST; argonaute-2, 1:1000, ab186733, Abcam; argonaute-2 (phospho Y393), 1:1000, ab215746, Abcam) and secondary HRP-conjugated IgG antibodies (1:5000, multi sciences, China). Bands were visualized by enhanced chemiluminescence and the band density was determined using ImageJ software.

### 2.9 Transfection

For each sample, a Lipofectamine2000 mixture was prepared as follows: dilute miRNA control /mimic/inhibitor (Gene Pharma., China): dilute with 50 μL of DMEM and 1.5 μL of a storage solution, then incubate at room temperature for 5 min; dilute liposome lipo2000: dilute 1 μL of lipo2000 with 50 μL of DMEM, incubate at room temperature for 5 min; mix, then incubate at room temperature for 20 min. The mixed solution was then added to the cells, and after 8 hours, the fresh medium was replaced. Transfection of point mutant plasmids: dilution of AGO-2 Y393F plasmid/AGO-2 WT plasmid (Gene Pharma., China), dilute with 50 μL of DMEM and 2.5 μL of plasmid, then incubate at room temperature for 5 min; diluttion of liposome lipo2000: dilute 7.5 μL of lipo2000 with 50 μL of DMEM, incubate at room temperature for 5 min; mix, then incubate at room temperature for 20 min. The mixed solution was then added to the cells, and after 8 hours, the fresh medium was replaced. The sequences of the RNA were as follows: Negative control: sense: 5’-UUC UCC GAA CGU GUC ACG UTT-3’, antisense: 5’-ACG UGA CAC GUU CGG AGA ATT-3’. miRNA-1-3p mimics: sense: 5’-UGG AAU GUA AAG AAG UAU GUA U-3’, antisense: 5’-ACA UAC UUC UUU ACA UUC CAU U-3’, miRNA-1-3p inhibitor: 5’-AUA CAU ACU UCU UUA CAU UCC A-3’, inhibitor NC: 5’-CAG UAC UUU UGU GUA GUA CAA-3’.

### 2.10 Statistical analysis

All the above experiments were performed at least three times, and data are calculated as mean ± SEM. GraphPad Prism Version 8 (GraphPad Software, San Diego, CA) software was utilized to determine statistical significance by unpaired Student’s t-test and one-way analysis of variance. The values of *P* < 0.05 were considered to be statistically significant.

## 3. Result

### 3.1 Chronic stress causes changes in microRNA levels in rat plasma and leads to increased level of coagulation-associated protein

After 4 weeks of modeling, rat microRNAs in plasma were down-regulated as determined by microarray (Figure 1A), and the most decreased microRNAs were microRNA-1-3p and microRNA-206-3p. It was verified by qPCR, results showed that the plasma levels of microRNA-1-3p were significantly decreased in the CUMS group compared with that in the control group, while microRNA206-3p showed a decreasing trend but no significant difference (Figure 1B). Next, we used Targetscan and miRDB to predict the target genes of these two microRNAs and found that they are all bound to the 3’UTR of TSP-1 (Figure 1C). The plasma levels of coagulation-related proteins platelet-associated factor (PAF), von Willebrand factor (vWF), and thrombospondin-1 (TSP-1) were measured by ELISA. Compared with the control group, there was no significant change in PAF and vWF, but the level of TSP-1 increased significantly. (Figure 2).

**Figure 1.**
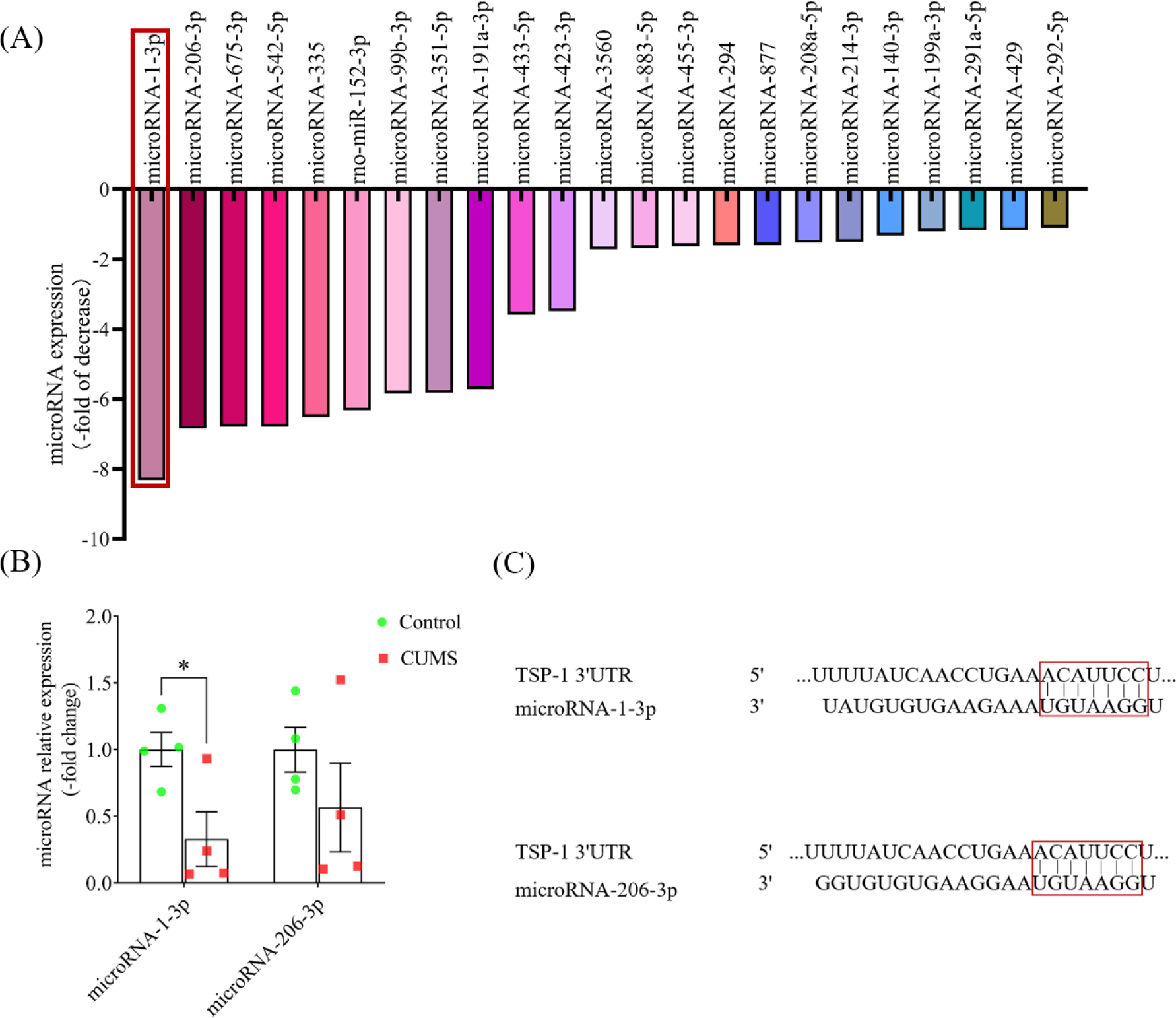
Chronic unpredictable mild stress (CUMS) induces changes in microRNAs in plasma. (A) Compared with the control group, the expression of microRNAs in the plasma of the stress group decreased. (B) The qPCR (Quantitative Real-Time PCR) was used to confirm the changes of microRNA-1-3p and microRNA-206-3p in plasma (n = 4), with microRNA-16 as the reference gene. (C) Bioinformatics tools (Targetscan and miRDB) were used to predict the potential targets of microRNA-1-3p and microRNA-206-3p. Data represent the mean ± SEM; \**P* < 0.05 *vs. Control* group.

**Figure 2.**
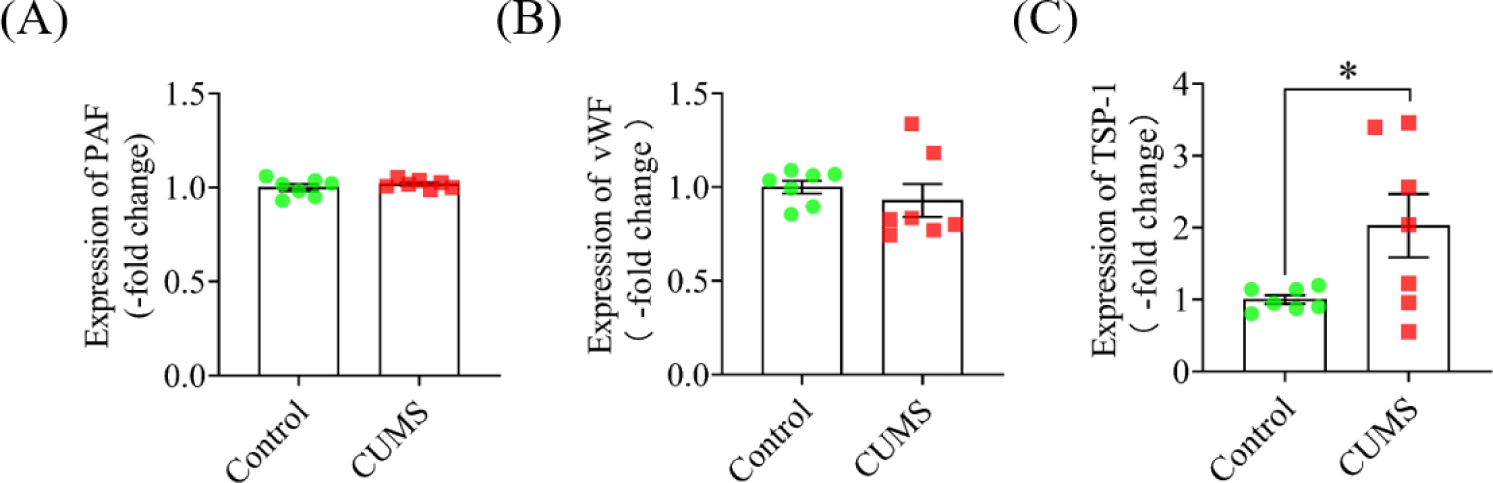
Chronic unpredictable mild stress (CUMS) causes changes in coagulation factors in plasma. (A) Plasma PAF levels were measured with ELISA. (B) Plasma vWF levels were measured with ELISA. (C) Plasma TSP-1 levels were measured with ELISA. Data were represented with the mean ± SEM (n=7); \**P* < 0.05 *vs. Control* group.

### 3.2 Glucocorticoid receptor activation leads to increased expression of TSP-1 in HUVECs

Stress leads to the release of cortisol (humans) and corticosterone (rodents), which in turn can lead to glucocorticoid receptor activation. We treated HUVECs with different concentrations of hydrocortisone sodium succinate (HSS) (0, 1, 3, 10, 30, and 100μM) for 6, 12, and 24 hours, extracted RNA when the cell density reached 80% to 90%, and detected changes in TSP-1 by qPCR (Figure 3). The increase of TSP-1 was most obvious at 10μM for 12 hours. Therefore, 10μM was selected as the administration concentration in subsequent experiments. We next verified by western blot and immunofluorescence that after treating the cells with 10μM HSS for 12 hours, the protein expression was significantly higher in the HSS group (Figure 4A and 4B) and the fluorescence intensity was also significantly higher (Figure 4C and 4D) compared to the control group. The increase of TSP-1 expression by HSS was completely blocked when the glucocorticoid receptor blocker RU-486 was pretreated (Figure 4E and 4F).

**Figure 3.**
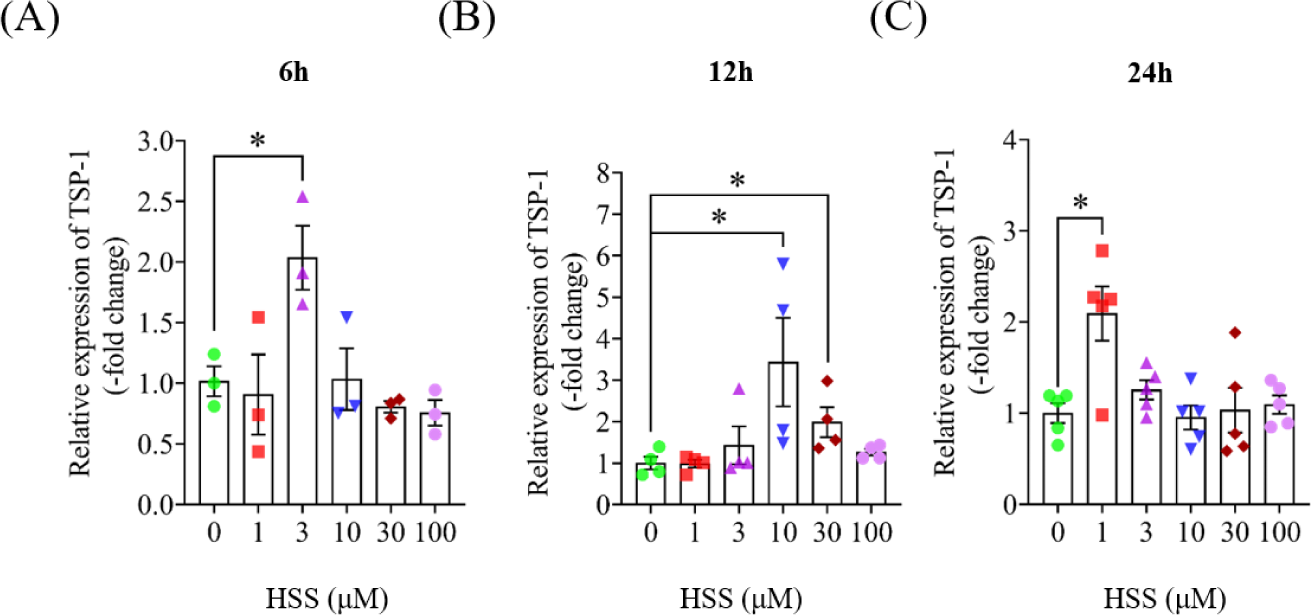
Effects of hydrocortisone sodium succinate (HSS) on expression of TSP-1 with different concentrations for a different time on TSP-1 in HUVECs. HUVECs grown to 70-80% in six-well plates were treated with 0, 1, 3, 10, 30, and 100 μM of hydrocortisone sodium succinate for 6, 12, and 24 hours, respectively, and the expression of TSP-1 was detected by qPCR. (A) The changes in TSP-1 mRNA content were detected after the HSS concentration gradient was applied to cells for 6 hours. (B) The changes in TSP-1 mRNA content were detected after the HSS concentration gradient was applied to cells for 12 hours. (C) The changes in TSP-1 mRNA content were detected after the HSS concentration gradient was applied to cells for 24 hours. Data represent the mean ± SEM (n = 3∼5); \**P* < 0.05 *vs. 0* group. The data were normalized to the expression of GAPDH.

**Figure 4.**
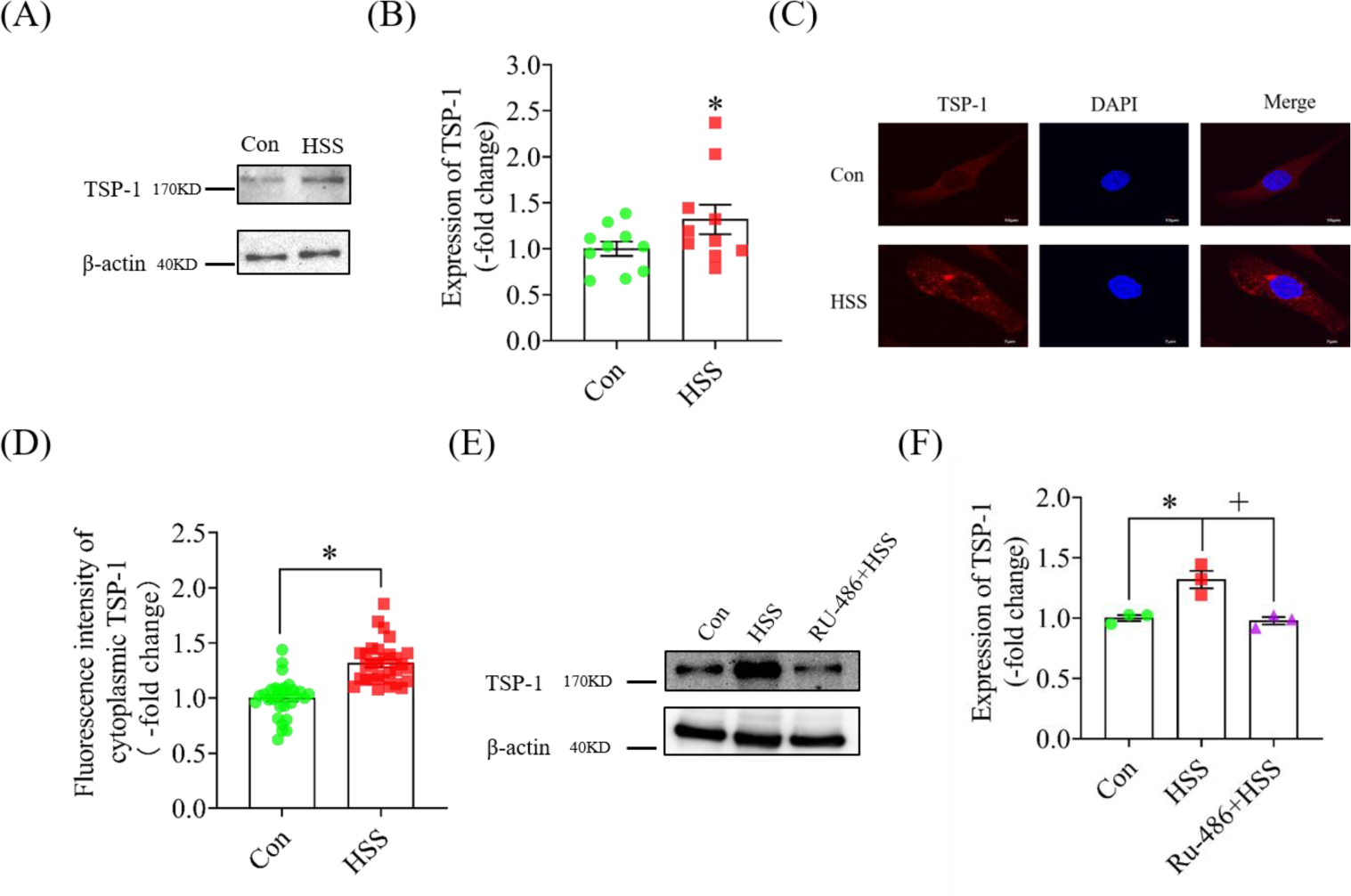
Glucocorticoid receptor activation upregulates TSP-1 expression. (A) Representative bands of western blot from HUVECs treated with HSS for 12 hours with β-actin as the internal reference. (B) Statistical graph corresponding to Figure (A) (n = 10). (C) Representative image of immunofluorescence. After TSP-1 immunofluorescence staining, TSP-1 was observed by laser confocal imaging. Fluorescence secondary antibody Alexa Fluor ®594 was used to label TSP-1, and DAPI was used to label nuclei. The scale bar = 7μm. (D) Statistics of the red fluorescence intensity of TSP-1 in Figure (C) (n =30). (E) Representative bands of western blot. RU-486 was pretreated for 30 min, and then HSS was added to treat HUVEC for 12 hours. β-actin was used as the internal reference. (F) Statistical graph corresponding to Figure (E) (n = 3). Data were represented the mean ± SEM; \**P* < 0.05 *vs. control* group, ^+^*P* < 0.05 *vs. HSS* group.

### 3.3 Glucocorticoid receptor activation leading to TSP-1 elevation is regulated through p38 MAPK and AGO-2 Y393 phosphorylation

HUVECs were treated with 10μM HSS for 10min, 30min, 1 hours, 3 hours, 6 hours, and 12 hours, and when the cell density in six-well plates reached 80%-90%, the protein was extracted and the phosphorylation of p38 MAPK was detected by western blot. The results showed a significant increase in the phosphorylation level of p38 MAPK at 3 hours compared to the control group (Figure 5A and 5B). The phosphorylation level of AGO-2 Y393 was also significantly increased at 3 hours and 6 hours (Figure 6A and 6B). When the p38 MAPK phosphorylation inhibitor birb796 was added, the phosphorylation levels of p38 MAPK (Figure 5C and 5D) and AGO-2 Y393 phosphorylation (Figure 6C - 6E) was reduced, as was the expression of TSP-1 (Figure 5E and F). In order to study the role of AGO-2 Y393 phosphorylation, we constructed the AGO-2 Y393F point mutant plasmid, transfected mutant plasmids into HUVECs, and detected the expression of TSP-1 by western blot. TSP-1 expression was decreased in the transfected Y393F group compared with the WT group (Figure 6F and 6G). Therefore, GR activation leads to elevated TSP-1 expression through phosphorylation of p38 MAPK /AGO-2.

**Figure 5.**
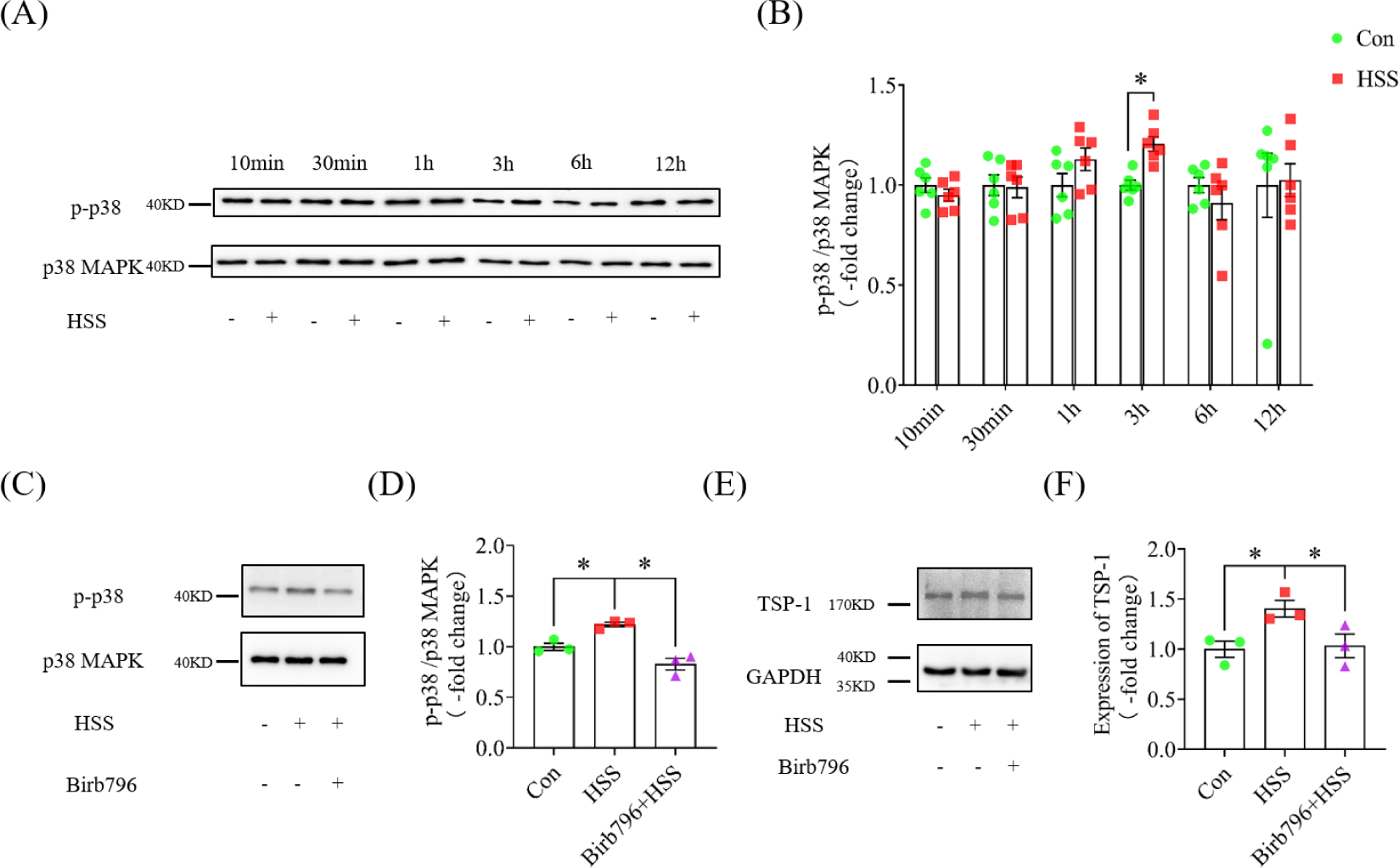
Phosphorylation of p38 MAPK plays an important role in this process. (A) Representative bands by western blot. Effect of HSS treatment of HUVECs at different times on p38 MAPK phosphorylation. (B) Statistical graph corresponding to Figure (A) (n = 6); \**P* < 0.05 *vs. Control* group. (C) Representative bands of phospho-p38 MAPK from western blot after inhibition of p38 MAPK phosphorylation. Effect of pretreatment with Birb796 for 2 hours and then HSS treatment of HUVECs 3 hours on p38 MAPK phosphorylation. (D) Statistical graph corresponding to Figure (C) (n = 3); \**P* < 0.05 *vs. HSS* group. (E) Representative bands of TSP-1 from western blot after inhibition of p38 MAPK phosphorylation. Effect of pretreatment with Birb796 for 2 hours and then adding HSS to treat HUVEC 12 hours on TSP-1. GAPDH was used as the internal reference. (F) Statistical graph corresponding to Figure (E) (n = 3); \**P* < 0.05 *vs. HSS* group. Data were represented the mean ± SEM.

**Figure 6.**
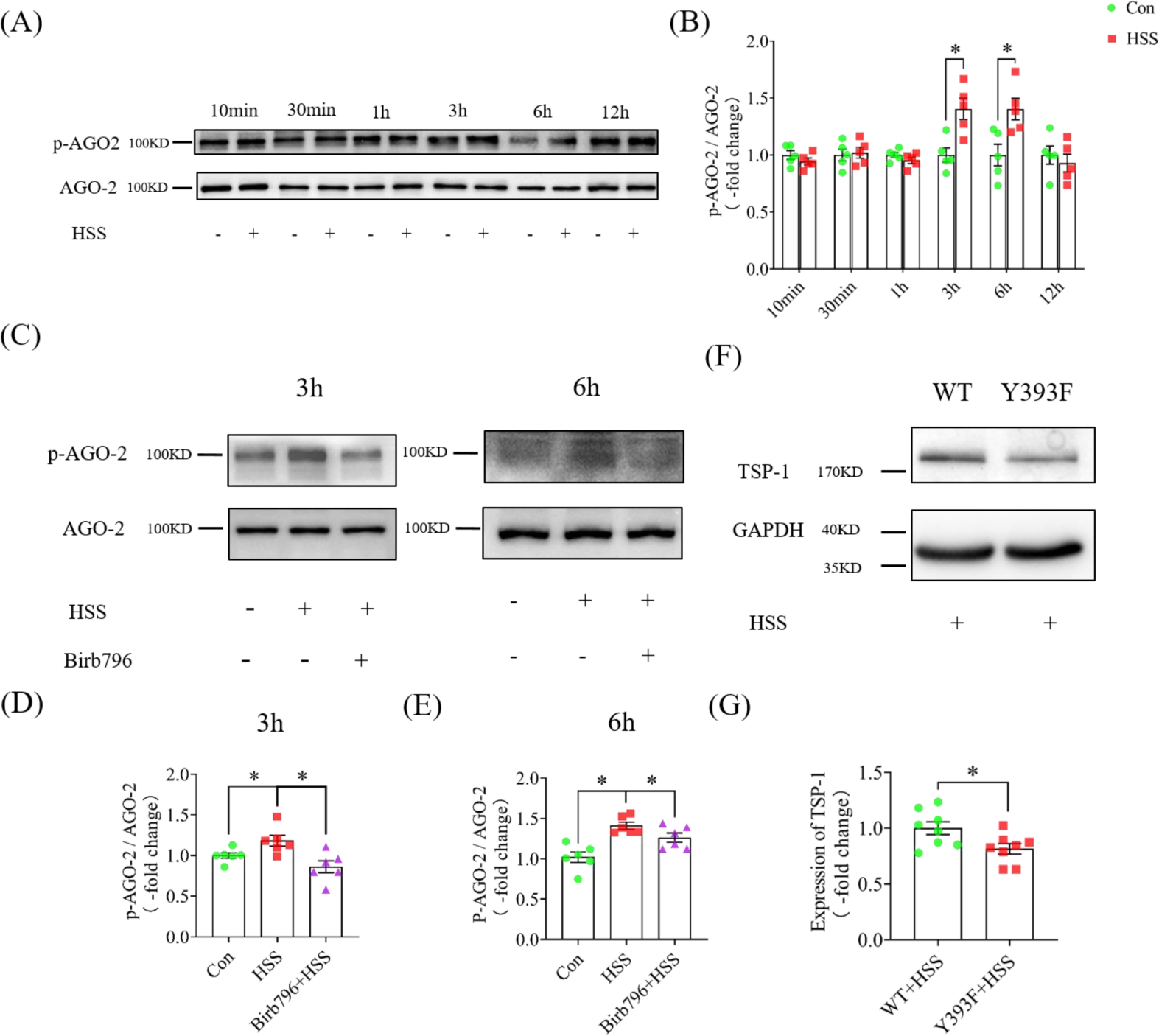
AGO-2 Y393 phosphorylation also plays an important role. (A) Effect of HSS treatment of HUVEC at different times on the phosphorylation of AGO-2 Y393. (B) Statistical graph corresponding to Figure (A) (n = 5); \**P* < 0.05 *vs. control* group. (C) Representative bands of phosphor-AGO-2 by western blot after treatment with p38 MAPK phosphorylation inhibitor. Birb796 was pretreated for 2 hours, and then HSS was added to treat HUVECs for 3 hours and 6 hours, respectively. Phosphorylation of AGO-2 Y393 was observed. (D) and (E) statistical graphs corresponding to 3 hours and 6 hours (n = 6); \**P* < 0.05 *vs. control* group. (F) Representative bands of TSP-1 by western blot after Y393F point mutation. Changes in TSP-1 were detected by western blot after transfection with AGO-2 WT plasmid and Y393F point mutant plasmid, respectively, GAPDH was used as the internal reference. (G) Statistical graph corresponding to Figure (F) (n = 8); \**P* < 0.05 *vs. wild+HSS* group. Data were represented the mean ± SEM.

### 3.4 MicroRNA-1-3p is also involved in regulating the expression of TSP-1

HUVECs were treated with 10μM HSS for 12 hours, microRNA was extracted, and qPCR showed that microRNA-1-3p content was significantly decreased compared with the control group, and the expression of microrNA-1-3p was increased after pretreated with birb796 (Figure 7A). Meanwhile, AGO-2 Y393 for phosphorylation can inhibit the maturation of microRNA-1-3p. We transfected the wild-type plasmid of AGO-2 and Y393F point mutant plasmid in HUVECs and detected the change of microRNA-1-3p by qPCR, the expression of microRNA-1-3p was significantly elevated in the Y393F group compared with the wild-type group (Figure 7B). To verify whether microRNA-1-3p targets the regulation of TSP-1, we transfected HUVECs with mimics or inhibitor of microRNA-1-3p and detected changes in TSP-1 by western blot. Results showed that transfection with microRNA-1-3p mimics decreased expression of TSP-1 (Figure 7C, 7D), while transfection with microRNA-1-3p inhibitor increased expression of TSP-1 (Figure 7E, 7F).

**Figure 7.**
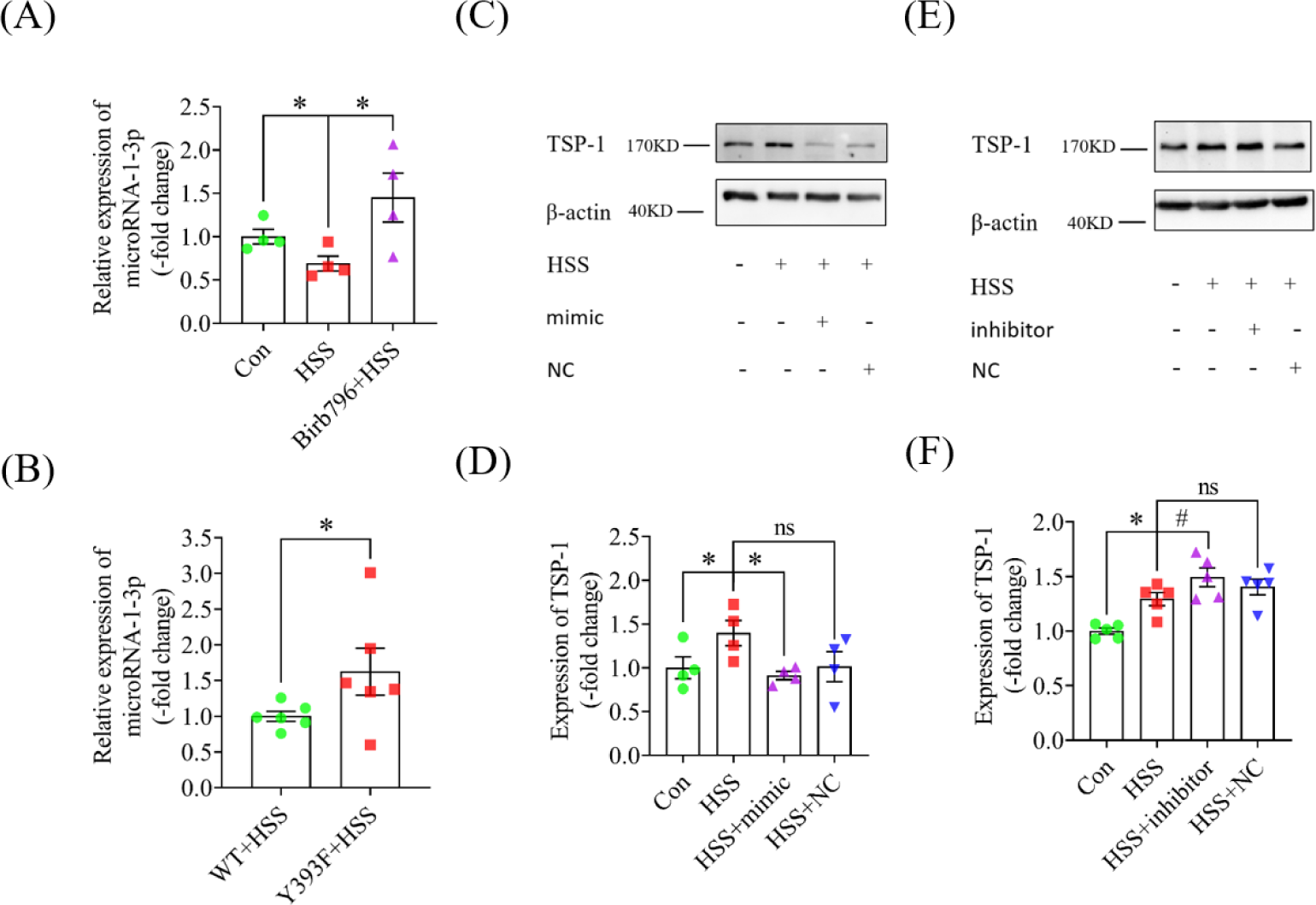
MicroRNA-1-3p is involved in the regulation of TSP-1 expression. (A) Birb796 was pretreated for 2 hours, HSS was added for 12 hours, microRNAs were extracted, and the changes of microRNA-1-3p were determined by qPCR. RNU6B was used as the internal reference (n = 4); \**P* < 0.05 *vs. HSS* group. (B) The AGO-2 wild-type plasmid and Y393F point mutation plasmid were transfected separately, and then HSS was added for 12 hours, microRNAs were extracted, and the changes of microRNA-1-3p were determined by qPCR. RNU6B was used as the internal reference (n = 6); \**P* < 0.05 *vs. wild+HSS* group. (C) The changes of TSP-1 after transfection with microRNA-1-3p mimics were detected by western blot. β-actin was used as the internal reference. (D) Statistical graph corresponding to Figure (C) (n = 4); \**P* < 0.05 *vs. HSS* group. (E) The changes of TSP-1 after transfection with microRNA-1-3p inhibitor were detected by western blot. β-actin was used as the internal reference. (F) Statistical graph corresponding to Figure (E) (n = 5); \**P* < 0.05 *vs. control* group, ^#^*P* < 0.05 *vs. HSS* group. Data were represented means ± SEM.

### 3.5 Glucocorticoid receptor activation in platelets leads to the release of TSP-1 and the increase of p38 MAPK phosphorylation

Platelets were obtained by differential centrifugation from rat, cultured in serum-free DMEM, and treated with 30 μM CORT for 0.5, 1, 3, 6 and 12 hours. The release of TSP-1 from the supernatant was detected by ELISA. Western blot was used to detect the phosphorylation of p38 MAPK and the expression of TSP-1. The results showed a significant increase of TSP-1 content in supernatant at 3 hours of GR activation (Figure 8A), indicating that GR activation leads to TSP-1 release from platelet α granules. Phosphorylation of p38 MAPK was also significantly increased at 12 hours of GR activation, whereas the expression of TSP-1 did not change significantly until 12 hours in the platelets (Figure 8B-8E). As shown in HUVECs, GR activation firstly leads to release of TSP-1 from platelets, in addition, it may also result in phosphorylation of p38 MAPK and finally to increase TSP-1 expression, probably because the treatment time is not long enough to reach the time point that causes changes in TSP-1 expression. However, since platelets cannot be cultured for a long time, they can only be cultured for 12 hours at most, which may need to be further verified by in vivo experiments.

**Figure 8.**
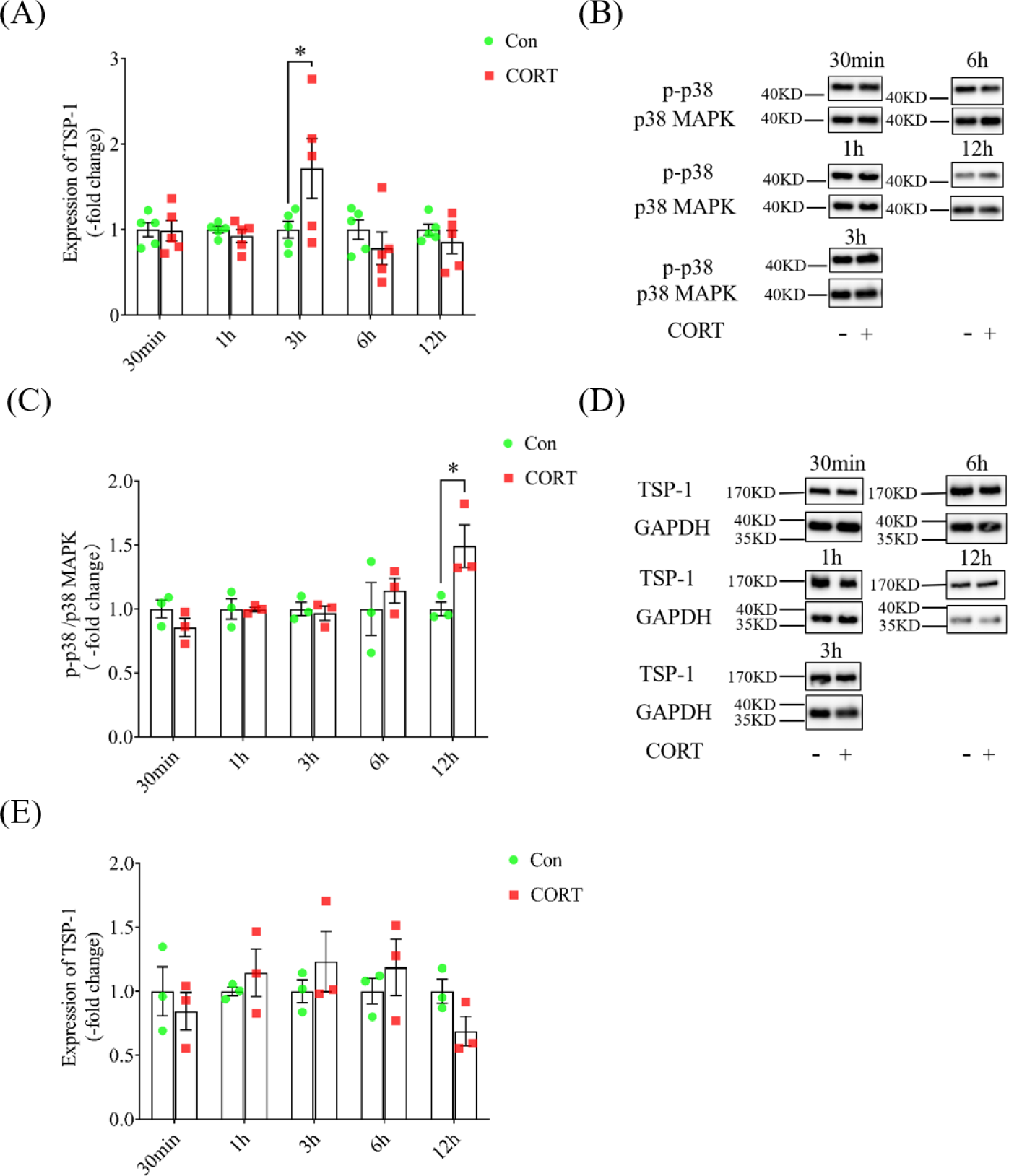
Glucocorticoid receptor activation promotes the release of TSP-1 from platelets and increase of p38 MAPK phosphorylation. (A) Platelets were treated with corticosterone (CORT) at different times, and the content of TSP-1 in cell supernatant was detected by ELISA (n = 5). (B) Platelets were treated with CORT at different times, and phosphorylation of p38 MAPK was detected by western blot. (C) Statistical graph corresponding to Figure (B) (n = 3). (D) Platelets were treated with CORT at different times. The changes of TSP-1 were detected by western blot. GAPDH was used as the internal reference. (E) Statistical graph corresponding to Figure (D) (n = 3). Data were represented the mean ± SEM; \**P* < 0.05 *vs. control* group.

## 4. Discussion

Earlier studies in our laboratory demonstrated that stress promotes platelet aggregation, leading to the exacerbation of deep vein thrombosis. So which pathway and which coagulation-related factor does it affect to promote platelet agglutination? We extracted serum from the abdominal aorta and examined the changes in PAF, vWF, and TSP-1, the results showed that the level of TSP-1 was significantly increased, while the levels of vWF and PAF were not significantly changed. vWF can mediate the adhesion of platelets to endothelial cells(52). It has been shown that oxidative stress can increase(53) and lead to endothelial damage(54) under stress conditions, and oxidative stress can up-regulate the expression of vWF in endothelial cells(55). vWF aggregates to form ultra-large vWF multimers. The conversion of vWF to vWF multimers in the blood is carried out by the VWF-specific metalloprotease ADAMTS-13, which cleaves the ultra-large vWF multimers into smaller, less adhesive multimers to prevent thrombosis(56). However, TSP-1 can competitively inhibit the binding of ADAMTS-13 to vWF and inhibit the cleavage of vWF by ADAMTS-13(57). We speculated that although the total serum content of vWF did not change, the increased content of TSP-1 might inhibit the cleavage of ADAMTS-13 to super large vWF, resulting in the formation of super large vWF, the ability to adhere to platelets was enhanced, platelet aggregation was promoted, and thrombosis was exacerbated. In addition, platelet aggregation is also regulated by some inhibitors, such as nitric oxide, prostaglandins and thrombin(58). It has been shown that stress can lead to endothelial injury and decrease the synthesis and release of nitric oxide(59). The enzyme plasminogen plasminogen is converted to plasmin by plasminogen activator to degrade fibrin within the clot to cause thrombolysis(60). It also has been documented that work stress leads to impaired fibrinolysis, including decreased tissue plasminogen activator and increased levels of plasminogen activator inhibitor(61). Therefore, we speculate that in addition to affecting the expression of procoagulation-related factors, chronic stress may also inhibit the expression of anticoagulant factors, leading to the imbalance of coagulation system homeostasis and promoting platelet aggregation, which needs to be further verified by subsequent experiments.

Previous studies have confirmed that chronic stress affects microRNA expression(29). Gene microarray analysis of microRNAs after RNA extraction showed significant differences in microRNA expression, with 47 microRNAs being significantly different, 24 being up-regulated and 23 being down-regulated. microRNA-1-3p and microRNA-206-3p were the most significantly decreased, and microRNA-1-3p was significantly decreased after being verified by qPCR. More research on microRNA-1-3p has focused on cancer. microRNA-1-3p is known to inhibit the proliferation of gastric cancer cells(62) and affect the resistance of lung cancer cells to gefitinib(63). However, whether microRNA-1-3p is involved in the effect of chronic stress on TSP-1 is still unclear. We predicted by bioinformatics software that microRNA-1-3p and microRNA-206-3p could bind complementary to the 3’UTR of TSP-1, indicating that these two microRNA may inhibit the translation of TSP-1 by binding complementary to the 3’UTR of TSP-1 in RISC. To test whether microRNA-1-3p can target TSP-1, we transfected microRNA-1-3p mimics and inhibitors to observe the effect of microRNA-1-3p on TSP-1. The results showed that when transfected with the mimic, the expression of TSP-1 was decreased, when transfected with the inhibitor, the expression of TSP-1 increased, indicating that microRNA-1-3p could target TSP-1 to inhibit the translation process of TSP-1. This suggests that microRNA-1-3p is involved in the regulation of TSP-1 expression in the process of chronic stress-induced changes in TSP-1. Since microRNA-206-3p also binds complementary to the 3’UTR of TSP-1, we speculate that microRNA-206-3p may also be involved in the regulation of TSP-1 expression during chronic stress-induced changes in TSP-1 expression. However, we did qPCR verification and found that microRNA-206-3p had a downward trend but no significant difference, so we focused on microRNA-1-3p.

Chronic stress rapidly activates the hypothalamic-pituitary-adrenal (HPA) axis to promote the synthesis and release of glucocorticoids(64). Glucocorticoids can bind to glucocorticoid receptors and activate glucocorticoid receptors. Our study demonstrated that GR activation regulates TSP-1 expression through microRNA-1-3p. MicroRNA biosynthesis is a multi-step process. The first step is the transcription of microRNA genes, mainly through RNA polymerase Ⅱ to form pri-microRNA^(65, 66)^. Pri-microRNA is further processed in the nucleus to form microRNA precursors with lenth of 70nt, also called pre-microRNAs. Pre-microRNA is transported from the nucleus and cleaved in the cytoplasm by Dicer, TAR RNA-binding protein (TRBP), and AGO-2 to form mature microRNA. AGO-2 is not only a component of RISC((67–69)), but also participates in microRNA maturation. A variety of post-translational modifications can modulate AGO-2 activity and function. It has been previously shown that phosphorylation of the AKT pathway at S387 alters the cellular localization of AGO-2 and promotes RISC activity(70). ERK-mediated phosphorylation of S387 enhances AGO-2 protein stability in neuronal cells(71) and prevents AGO-2 secretion into exosomes(72). Phosphorylation of Y529 reduces the localization of AGO-2 P-bodies(73) while affecting microRNA binding to AGO-2(74). EGFR interacts with AGO-2 under hypoxic conditions, leading to increased phosphorylation of Y393 and hindering the binding of AGO-2 to dicer to inhibit the microRNA maturation process(47). Our experiments demonstrated that glucocorticoid receptor activation promoted p38 MAPK phosphorylation at 3 hours, AGO-2 Y393 phosphorylation at 3 hours and 6 hours, and decreased microRNA-1-3p content. The phosphorylation of AGO-2 Y393 may also affect the binding of AGO-2 to Dicer and affects the maturation of microRNA. When we mutate the tyrosine of AGO2 393 to phenylalanine, AGO-2 393 cannot be phosphorylated, and the content of microrNA-1-3p is reduced. Our study for the first time demonstrated the role and mechanism of microRNA-1-3p and its regulation on TSP-1 in enhanced platelet aggregation induced by chronic stress, as well as the mechanism by which chronic stress regulates microRNA-1-3p.

Our study demonstrated that p38 MAPK phosphorylation and AGO-2 phosphorylation regulated the maturation of microRNA-1-3p and hence regulated TSP-1 expression after GR activation in HUVECs. TSP-1 is expressed in cells such as megakaryocytes, macrophages, and smooth muscle cells(15), while glucocorticoid receptors are almost ubiquitous and have been confirmed to be expressed in smooth muscle cells, megakaryocytes, and macrophages((75–77)), In addition, TSP-1 is largely stored in platelet α granules(14). Although platelets are anucleated, they can regulate the transcription of mRNA (inherited from the megakaryocytes from which they are derived)(78), so we speculate that platelet release of TSP-1 upon glucocorticoid receptor activation would regulate TSP-1 expression by the same mechanism as in HUVEC. Our results showed that GR activation led to the release of TSP-1 and the phosphorylation of p38 MAPK, but the protein expression of TSP-1 did not change significantly. At present, platelet culture in vitro can only be cultured for 12 hours, so we speculate that the culture time may not be enough, and we need to do in vivo experiments to verify.

In conclusion, our study demonstrates that GR activation in HUVEC stimulates the phosphorylation of p38 MAPK, which in turn promotes the phosphorylation of AGO-2 and inhibits the maturation of microRNA-1-3p, leading to elevated expression of TSP-1, GR activation in platelets leads to the release of TSP-1. Our observation may provide novel strategy in clinic to prevent chronic stress induced stroke, cardiac infarction and development of thrombosis.

## Funding

This work is supported by the National Natural Science Foundation of China (81970422), Suzhou Municipal Science and Technology Bureau (SS201745, SYS2019108, GSWS2019022), Suzhou Commission of Health (GSWS2019022) and Jiangsu Commission of Health (LGY2019012).

The authors would like to thank all the reviewers who participated in the review and MJ Editor (www.mjeditor.com) for its linguistic assistance during the preparation of this manuscript.

